# Rapid, high-yield production of full-length SARS-CoV-2 spike ectodomain by transient gene expression in CHO cells

**DOI:** 10.1101/2020.09.08.286732

**Authors:** Matthew Stuible, Christian Gervais, Simon Lord-Dufour, Sylvie Perret, Denis L’Abbe, Joseph Schrag, Gilles St-Laurent, Yves Durocher

## Abstract

Recombinant forms of the spike protein of SARS-CoV-2 and related viruses have proven difficult to produce with good yields in mammalian cells. Given the panoply of potential COVID-19 diagnostic tools and therapeutic candidates that require purified spike protein and its importance for ongoing SARS-CoV-2 research, we have explored new approaches for spike production and purification. Three transient gene expression methods based on PEI-mediated transfection of CHO or HEK293 cells in suspension culture in chemically-defined media were compared for rapid production of full-length SARS-CoV-2 ectodomain. A high-cell-density protocol using DXB11-derived CHO^BRI/rcTA^ cells gave substantially better yields than the other methods. Different forms of the spike were expressed, including the wild-type SARS-CoV-2 sequence and a mutated/stabilized form (to favor expression of the full-length spike in prefusion conformation), with and without fusion to putative trimerization domains. An efficient two-step affinity purification method was also developed. Ultimately, we have been able to produce highly homogenous preparations of full-length spike, both monomeric and trimeric, with yields of 100-150 mg/L. The speed and productivity of this method support further development of CHO-based approaches for recombinant spike protein manufacturing.

## 1. INTRODUCTION

The availability of recombinant forms of the SARS-CoV-2 spike protein is becoming increasingly important for development of novel strategies to combat the current COVID-19 pandemic. In the human immune response to the SARS coronavirus, the spike protein is a key antigen and is often the target of neutralizing antibodies found in convalescent patients[1]. Recombinant spike protein has a wide range of potential applications, including in the development of novel therapeutics (e.g. neutralizing antibodies or other spike-targeting biologics), clinical diagnostic tools (e.g. to evaluate post-infection immunity) or subunit vaccines (different forms of recombinant SARS spike protein were effective as vaccines in animal models) [2, 3].

The SARS-CoV-2 spike protein is closely related to that of SARS-CoV-1, the virus responsible for the 2003 SARS outbreak. Both proteins are large, multi-domain glycoproteins with transmembrane domains that traverse the viral envelope and are proteolytically processed into S1 and S2 subunits. Notably, while the SARS-CoV-1 spike is only cleaved during infection of target cells, the SARS-CoV-2 spike contains a conserved furin recognition site at the S1/S2 junction, such that cleavage occurs during biosynthesis in host cells; this difference may impact the route of entry of the two virus types into host cells [4]. The SARS-CoV-1 spike protein was shown to assemble into homo-trimeric complexes that are found on mature viral particles [5]. When expressed in the absence of its transmembrane and C-terminal domains, the spike ectodomain (ECD) is reported to be mostly monomeric; fusion of the spike ECD C-terminus to the trimerization domain of bacteriophage T4 fibritin (T4-Fib or foldon) increases its capacity to elicit neutralizing antibodies [6].

In the literature on SARS-CoV-1 and other related coronaviruses, there are reports of various approaches for producing recombinant spike proteins. Individual domains of the spike, including the receptor-binding and hemagglutinin-esterase domains, have been produced in CHO, HEK293, Vero and insect cells [6-8]. It is also possible to express the full-length spike, including transmembrane and C-terminal domains, which can be purified following membrane solubilisation of expressing cells [9, 10]. Finally, expression of the full-length soluble forms of the coronavirus spike ECDs has also been reported in HEK293 and insect cells [6, 11-15]. Importantly, however, in cases where this data is reported, yields were extremely low, ranging from 0.5-1.5 mg per litre of culture media (for expression of constructs containing the full-length ECD) [9, 14]. More recently, for the SARS-CoV-2 spike, higher titers have been achieved using a highly engineered, stabilized trimeric spike ECD variant, reaching up to 33 mg/L by transient gene expression (TGE) in CHO [16]. Another recent paper used optimized processes and expression vectors to obtain yields of trimeric spike ECD of up to 10 and 6 mg/L by TGE in 293 and CHO, respectively [17]. Using methionine sulfoximine (MSX) selection to establish a stable CHO pool, the same authors reported a further increase in titers of up to 53 mg/L. While significant improvements have recently been achieved, these productivities are well below desirable levels for mass production, in particular for development and manufacturing for potential diagnostic or vaccine applications at commercial-scale.

Recently, we have attempted to produce the full-length SARS-CoV-2 spike ECD using three rapid protein production platforms established previously in our group, based on polyethylenimine (PEI)-mediated plasmid transfection of CHO or HEK293 cells. Yields of soluble ECD from the two CHO-based methods were substantially higher than other reported methods, with one method yielding approximately 100-150 mg/l (depending on the form of the spike expressed) at 6-7 days post-transfection. Both the wild-type spike and a stabilized form with mutated protease cleavage sites [14] were produced and purified; both forms were expressed with and without fusion to various putative trimerization domains. The CHO-secreted wild-type protein is a mix of unprocessed precursor and cleaved forms, likely including free S1, S2 and S1/S2 complexes. The stabilized protein (with protease site mutated) with or without trimerization domain fusion is secreted in a full-length, unprocessed form. Notably, for the wild-type (non-stabilized, furin site present) spike ECD constructs, we have observed dissociation of S1 from the monomeric and trimeric S1/S2 complexes during purification and storage, indicating that the furin site-deleted mutant is a better choice for ongoing production of the full-length ECD. The yield and purity we have obtained support continued method development to further improve titers and downstream process for eventual mass production in CHO cells.

## 2. MATERIALS & METHODS

### 2.1 Expression vectors

SARS-CoV-2 spike protein sequences were derived from Genbank accession number MN908947. Protein-coding DNA sequences were codon-optimized for CHO cells, synthesized and cloned into the pTT5™ vector [18] by Genscript. The specific regions of the spike ECD contained in each construct and the sequences and locations of the stabilizing mutations in ECDm are indicated in Figure 1. Trimerization domains were fused to the C-termini of the spike protein coding sequences followed by FLAG and 6xHis tags. Two constructs (pTT5-ECD-T4Fib and pTT5-ECDm-T4Fib) contain additional dual StrepTag-II tags between the FLAG and 6xHis tags.

**Figure 1:**
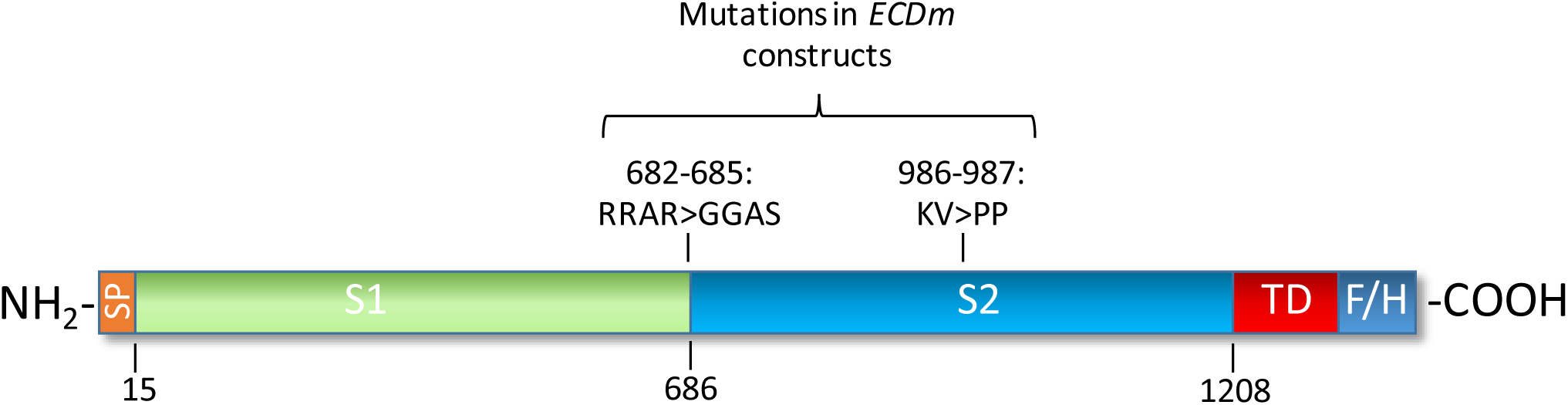
Organization of SARS-CoV-2 spike protein constructs. Full-length ectodomain (ECD) constructs consist of the native SARS-CoV-2 spike N-terminal signal peptide (SP) and remaining sequence up to amino acid 1208, including full S1 and S2 domains (native transmembrane and C-terminal domains were removed). Mutations present in the prefusion-stabilized (K986P, V987P), furin site-mutated (residues 682-685 [RRAR] mutated to GGAS) ECD (ECDm) are shown above. Trimerization domains (TD) and FLAG/His (F/H) affinity tags were introduced after the C-terminus of S2.

### 2.2 Expression platforms

Transfection and post-transfection culture of CHO-3E7 and 293-6E cells in FreeStyle F17 media were performed as described previously [19, 20].

The method for transient expression in CHO^BRI/rcTA^ cells [21] is based on a published method for high-density transfection of CHO-3E7 cells [19], with several modifications. Cells were maintained in shake flasks in a humidified, 5% CO_2_, 37°C incubator, rotating at 120 rpm in a chemically-defined proprietary media formulation. Cells were seeded in the same media at 1×10^6^ cells/ml 2 days prior to transfection to achieve a cell density of ∼8×10^6^/ml at the time of transfection. Before transfection, cells were diluted with 25% fresh media and dimethylacetamide was added to 0.083% (v/v). Transfections were performed by adding PEI-DNA polyplexes (10% final culture volume) to diluted cells (90% final culture volume). To prepare polyplexes, PEI-Max (Polysciences) and plasmid DNA were diluted separately to 200 ug/ml and 28 ug/ml, respectively, in a volume of growth media equal to 5% of the final culture volume. Plasmid DNA consisted of 85% of the different pTT5™-spike constructs, 10% pTT™-Bcl-XL (anti-apoptotic effector) and 5% pTT™-GFP. The diluted PEI-Max was added to the diluted DNA and incubated for 7 minutes at room temperature before adding to cells. At 24h post-transfection, cultures were shifted to 32°C and supplemented with Anti-Clumping Supplement (1:500 dilution) and Feed 4 (2.5% v/v), both from Irvine Scientific. Additional Feed 4 (5%) was added at 5 days post-transfection. Glucose concentrations were monitored every 2-3 days to maintain >10 mM. Cell supernatants were harvested at 6-7 days post-transfection (except for time-course experiments shown in Figure 3 and Supplementary Figure 1).

**Figure 2:**
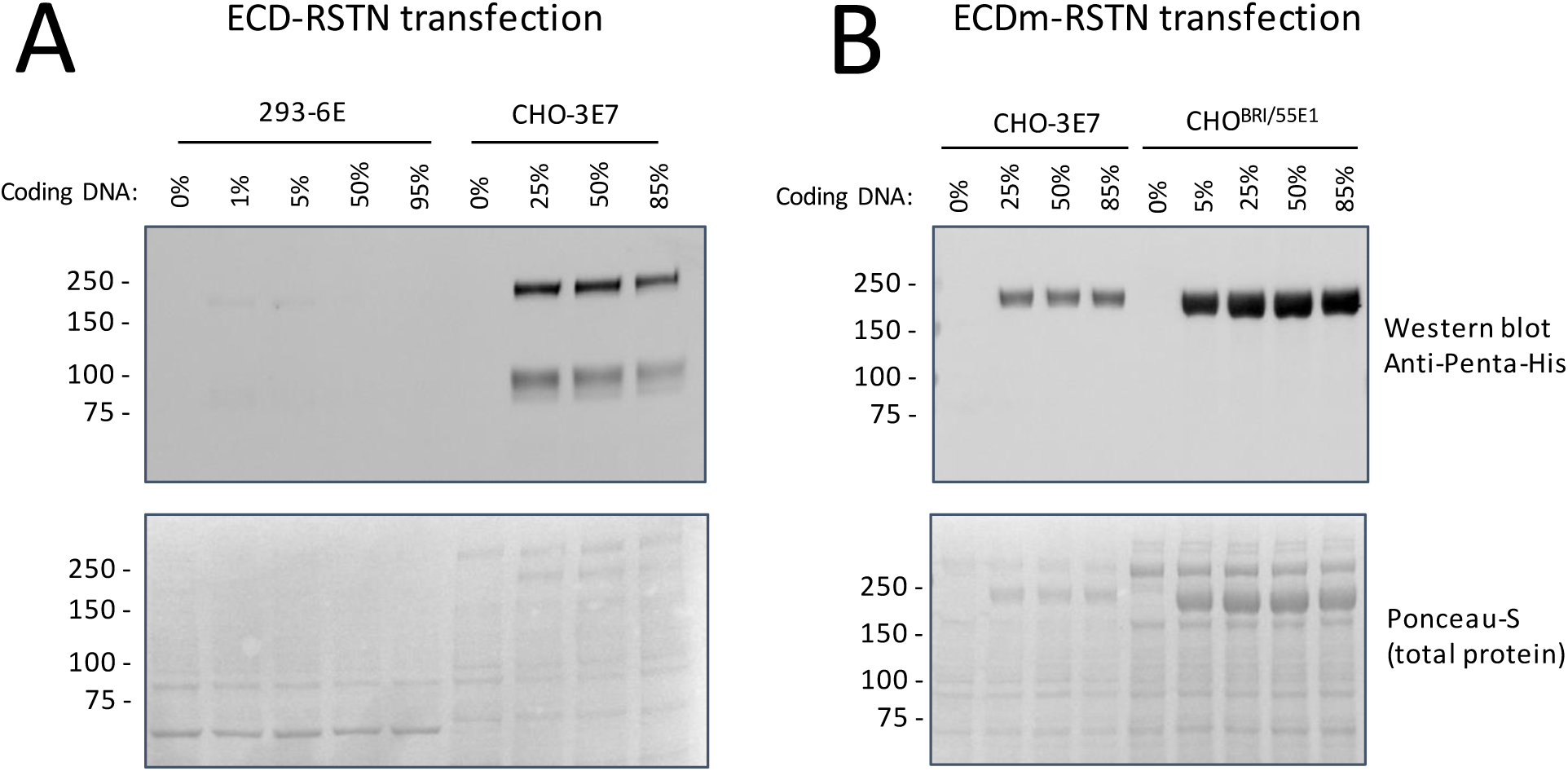
Evaluation of mammalian cell expression platforms for expression of SARS-CoV-2 ectodomain. 293-6E, CHO-3E7, or CHO^BRI/55E1^ cells were transfected with the indicated percentages of ECD-coding plasmid DNA. (A) Comparison of ECD-RSTN production in 293-6E and CHO-3E7 cells. Samples of culture supernatant were taken at 5 days post-transfection (dpt). (B) Comparison of ECDm-RSTN production in CHO-3E7 and CHO^BRI/55E1^ cells. Samples of culture supernatant were taken at At 5 dpt (CHO-3E7) or 6 dpt (CHO^BRI/55E1^). For all samples, 10 μl of culture supernatant was analyzed by western blotting. Ponceau-S staining of membranes before western blotting is shown in lower panels.

**Figure 3:**
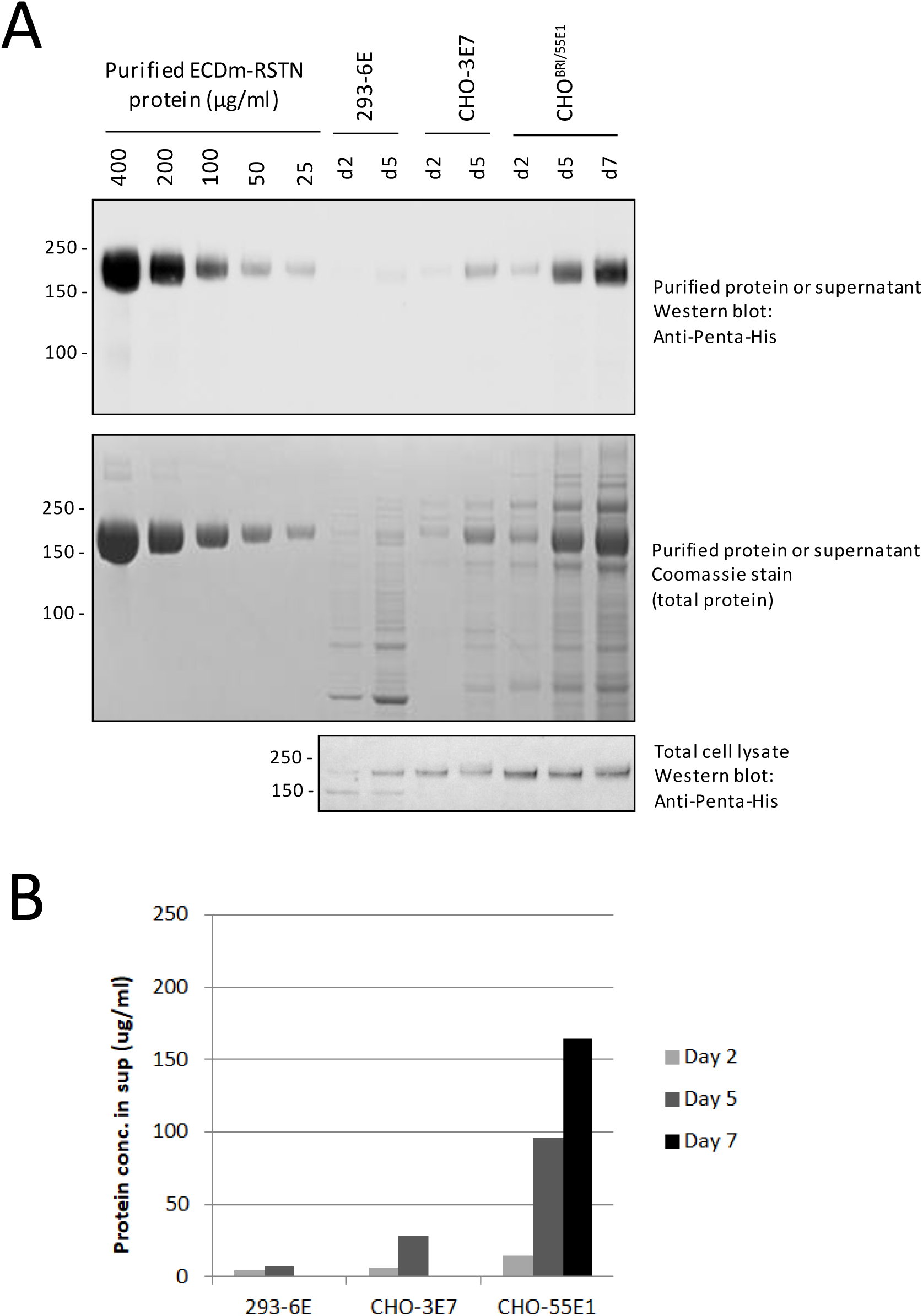
Evaluation of recombinant ECDm-RSTN concentrations in culture supernatants from transfected CHO-3E7, CHO^BRI/55E1^ and 293-6E cells. Cells were transfected with the ECDm expression plasmid. Samples were taken at 2, 5 or 7 days post-transfection for preparation of cleared culture supernatants and cell pellets (for lysis and total protein extraction). (A) Cleared supernatants (10 μl) and total protein extracts (equal amounts of total protein for all samples) were analyzed by western blotting along with known amounts of IMAC/FLAG-purified ECDm-RSTN protein. Densitometry analysis of the purified protein bands was used to prepare a standard curve and estimate ECD protein concentrations in the supernatants (B).

SDS-PAGE was performed using NuPAGE 4-12% Bis-Tris gels (Invitrogen). Total protein staining (Coomassie Blue) and western blotting were performed using standard methods. Anti-Penta-His Alexa Fluor 488 conjugate was from Qiagen.

### 2.3 Purification

All protein constructs were initially purified by IMAC on Nickel Sepharose Excel resin (GE Healthcare). 0.2 µm-filtered supernatants were applied to columns by gravity flow. The column volume used for a given volume of supernatant was variable for different productions; for maximum recovery of ECD constructs from cell supernatant, a column volume of 1 ml per 0.5 mg of protein is optimal. Columns were washed with 50 mM sodium phosphate, pH 7.0, containing 300 mM NaCl and 25 mM imidazole and eluted with the same buffer containing 300 mM imidazole.

In order to achieve near homogeneity, a second step purification was required. For most constructs, ANTI-FLAG M2 affinity gel (Sigma Aldrich) was used in batch mode. IMAC-purified proteins were incubated with the gel for ∼ 2 hours with gentle overhead mixing. One cc of gel was used for every 0.5 mg of IMAC-purified protein. Resin was collected by filtration in an empty column (Qiagen) and washed 3 times with 10 x CV of calcium- and magnesium-free Dulbecco’s PBS (DPBS). Bound proteins were eluted by competition with four 2 x CV of DPBS containing 100 µg/mL FLAG peptide (Sigma Aldrich). Purified proteins were formulated in DPBS by buffer exchange through desalting columns (GE Healthcare), 0.2 µm filtered (Millipore) and stored at -80°C at protein concentration not exceeding 2 mg/ml.

For constructs containing the T4-Fib trimerization domain, a StrepTactin column was used for the second purification step instead of anti-FLAG. IMAC eluate was loaded on a 5-ml StrepTrap HP column (Cytiva) at a flow rate of 5 ml/minute. The column was washed with 5 column volumes of and bound protein was eluted with DPBS containing 2.5 mM desthiobiotin. Buffer exchange into DPBS was performed as described above.

### 2.4 UPLC-SEC/MALS

UPLC-SEC-MALS analyses were conducted on a 4.6 × 150 mm BEH200 SEC column with 1.7 µm particles (Waters, Milford, MA) connected to an Acquity H-Class Bio UPLC system (Waters) with a miniDAWN™ MALS detector and Optilab^®^ T-rEX™ refractometer (Wyatt Technology, Santa Barbara, CA, USA). The column temperature was 30°C and the mobile phase was DPBS (HyClone SH30028.01) with 0.02% Tween 20 added. The flow rate was 0.4 mL/min. Weighted average molecular mass (*M*_MALS_) was calculated in ASTRA 6.1 software (Wyatt) using a protein concentration determined from the refractive index signal with a dn/dc value of 0.185.

## 3. RESULTS AND DISCUSSION

For protein expression, we used plasmid vectors containing CHO codon-optimized sequences encoding the full ECD of the SARS-CoV-2 spike protein. C-terminal FLAG and 6xHis tags were added for purification. Two versions of the protein were expressed: one encoding an unmodified ECD protein sequence (ECD) and a second with two modifications, to block furin-mediated S1/S2 cleavage and to stabilize the pre-fusion conformation (ECDm), as recently described [14]. To attempt to mimic the native trimeric structure of the spike protein, as found on the virus surface, trimerization domains of T4-Fibritin (T4-Fib) [6], human resistin (RSTN) [22], or GCN4 [23] were fused to the C-terminus of the ECD in certain constructs. A fourth, proprietary trimerization sequence (T3) was also tested.

We rapidly assessed the potential of three expression platforms for production of SARS-CoV-2 spike: two methods, based on EBNA1-expressing CHO (CHO-3E7) [19] and HEK293 (293-6E) [20] have served as core platforms for recombinant protein production in our group for several years, and generally perform very well for a wide range of recombinant antibodies and other proteins. For both methods, cells are cultured in chemically-defined F17 media and transfected at low cell density using PEI. A third method uses a CHO-DXB11-derived clone (CHO^BRI/RCTA^) that expresses machinery for cumate-inducible protein expression [21]; CHO^BRI/RCTA^ cells are used by our group for stable cell line development for biologics manufacturing. Recently, we have developed a method for high-cell-density PEI-mediated transfection of these cells which performs well with both constitutive and cumate-inducible promoters (unpublished data).

We began by transfecting CHO-3E7 and 293-6E cells with various amounts plasmid DNA encoding the spike ECD fused to the resistin trimerization domain (ECD-RSTN). Expression levels were compared at 5 days post-transfection; for other proteins, we frequently observe that while keeping the total amount of transfected DNA constant, reducing the amount of coding vs. non-coding (salmon sperm) DNA can improve yields [20]. As shown in Figure 2A, western blotting of bulk supernatants with an anti-penta-His antibody shows two clear bands in the CHO-3E7 supernatants transfected with 25-85% coding DNA, but little signal in the 293-6E supernatants transfected with 1-95% coding DNA. Two bands are detected in the anti-penta-His blots, at ∼80 kDa and ∼180 kDa, which likely correspond to the S2 subunit (after cleavage at the S1/S2 site) and the unprocessed precursor, respectively. These two proteins can be observed as faint bands by Ponceau S staining of membranes prior to western blotting. Based on amino acid sequence, the predicted molecular weight of the unprocessed spike is 143 kDa. The higher observed molecular weight is very likely due to glycosylation: the SARS-CoV-2 spike protein contains 22 confirmed N-linked glycosylation sites [24]. In addition, for the SARS-CoV-1 spike, enzymatic removal of glycans decreases apparent molecular weight by SDS-PAGE by 30-40 kDa [25].

Similar expression tests were performed for the stabilized spike protein (ECDm-RSTN) with the CHO-3E7 and CHO^BRI/RCTA^ platforms, using a range of coding DNA ratios. As shown in Figure 2B, an anti-penta-His western blot using unpurified supernatants gave a single band at ∼180 kDa, which corresponds to the size of the unprocessed precursor protein observed with expression of the non-stabilized spike (Figure 2A). The CHO^BRI/RCTA^ method gives substantially higher yields, based on band intensity on the anti-penta-His blot and Ponceau S stained membrane. For both cell lines, there was little difference in expression between 25 and 85% of coding DNA.

To estimate the concentration of spike protein in the culture supernatants, we performed western blots with unpurified supernatants from 293-6E, CHO-3E7 and CHO^BRI/RCTA^ cells transfected with ECDm-RSTN along with a series of dilutions of purified ECDm protein (Figure 3). Samples were taken up to 7 days post-transfection for CHO^BRI/RCTA^ cells and up to 5 days post-transfection for CHO-3E7 and 293-6E cells (CHO-3E7 and 293-6E cell viability dropped to <70% on Day 5). As shown in Figure 3B, densitometry analysis of this blot allows us to estimate that yields in the transfected CHO^BRI/RCTA^ supernatants reach >150 μg/ml at 7 days post-transfection. As expected, concentrations in CHO-3E7 supernatants were lower (∼25 μg/ml at Day 5) while they were <10 μg/ml in 293-6E supernatants. A similar comparison was performed for the ECDm protein without trimerization domain in CHO^BRI/RCTA^ and 293-6E cells (Supplementary Figure 1); in this case CHO^BRI/RCTA^ yields were ∼100 μg/ml and 293-6E yields were ∼20 μg/ml.

The differences in yields from the three cell lines were reproducible and were not due to differences transfection efficiency: for all transfections, we either co-transfected 5% of a GFP expression vector or included a GFP-only transfected external control, and we monitored GFP expression at 24-48 hours post-transfection (data not shown). Also, we did not observe any obvious negative effect of spike protein expression on cell growth or viability.

The differences in the amounts of spike protein secreted by the three cell lines are not reflected by intracellular protein levels. As shown in Figure 3A and Supplementary Figure 1C, recombinant spike protein is easily detectable in total protein extracts from all three cell lines by western blotting. Intracellular protein levels are slightly lower in CHO-3E7 and 293-6E cells compared to CHO^BRI/RCTA^ in Figure 3A, but certainly, these differences do not reflect the substantial differences in concentrations of secreted protein. These results confirm that all three cell lines were transfected and appear to be able to produce intracellular spike protein at similar levels; however, an explanation for the low level of protein secretion, especially in 293-6E cells, is still elusive.

Based on these results, we decided to pursue the CHO^BRI/RCTA^ TGE platform for ECD expression, and we proceeded to develop a purification method for supernatants from these cells. Bulk supernatants were first passed on an IMAC (nickel-Sepharose) column; as shown in Figure 4 for the ECD-T3 construct, the spike protein is prominent in the eluate from this column; as expected, three forms of the protein are observed: full-length S1/S2 as well as the processed S1 and S2 subunits. Several other non-specific proteins are also present. Following a second-step purification of this eluate on an anti-FLAG column, these contaminating proteins were effectively removed. Importantly, we observed a significant amount of S1 subunit in the flow-through and washes from the anti-FLAG column for the ECD-T3 construct as well as all other constructs with intact protease cleavage sites. We believe that this represents slow dissociation of S1 from S2 (the FLAG/His tag is fused to S2); S1 does not immediately dissociate from S2 after S1/S2 processing, but it is evident over time with multiple steps of purification/storage. Certainly, this result supports the use of the mutated form of the spike ECD (ECDm), lacking the S1/S2 cleavage site, for production of full-length recombinant spike protein.

**Figure 4:**
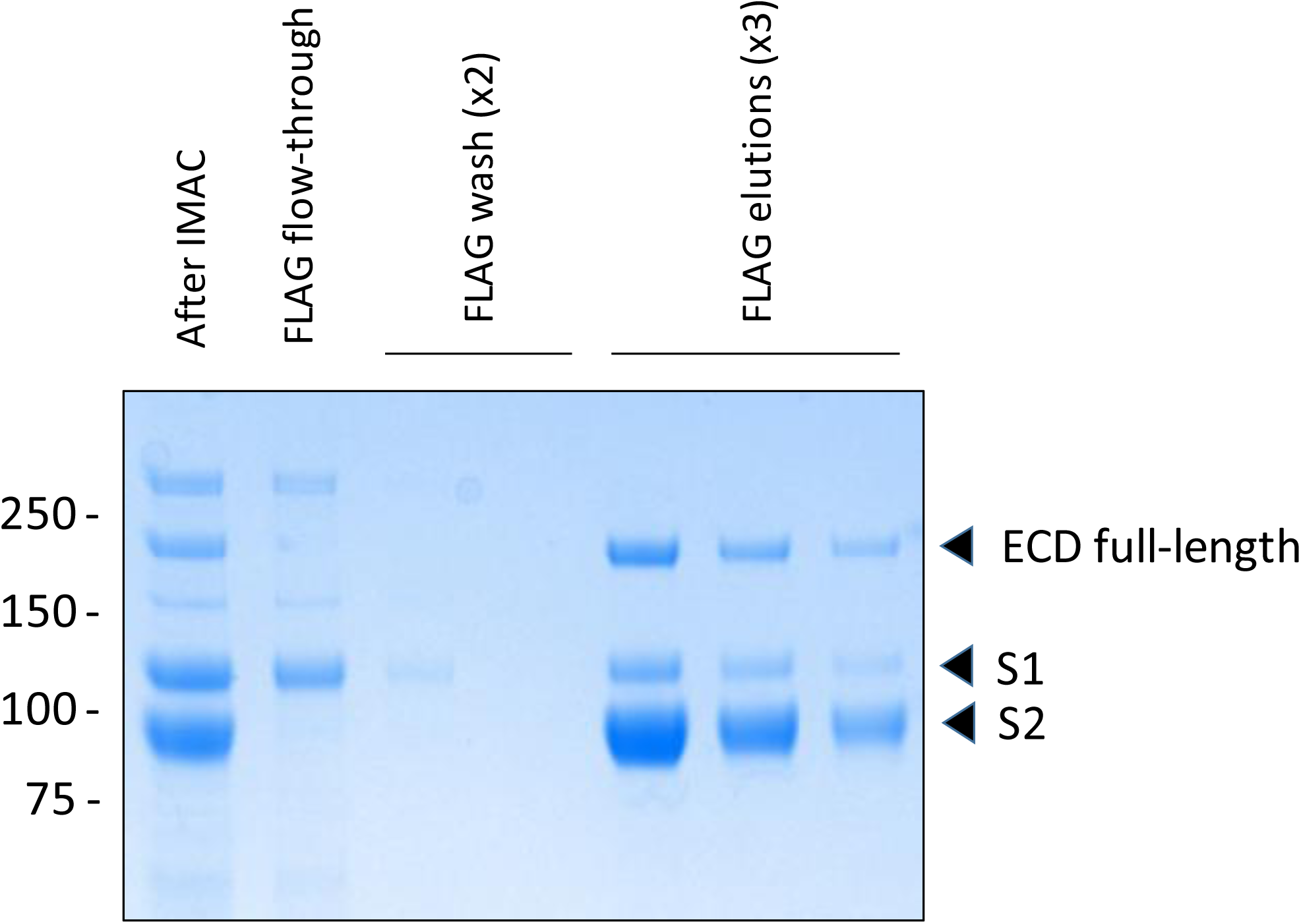
Loss of S1 subunit during second-step FLAG purification of ECD-T3. Proteins eluted from IMAC column (After IMAC) were purified with anti-FLAG beads. Flow-through, washes and elutions from FLAG column were analyzed by SDS-PAGE and Coomassie Blue staining.

Our purification method has evolved with our growing experience purifying different spike ECD constructs. In particular, in our initial IMAC purification runs, a large proportion (∼50%) of the protein was found in the column flow-through. The standard binding capacity for the Ni-Sepharose excel IMAC column is >10 mg/ml, but after considerable effort troubleshooting this step, we have concluded that the capacity is much lower for the spike ECD proteins (∼0.5 mg/ml). With an appropriate amount of supernatant loaded on the IMAC column, we are now able to capture the expected amount of ECD protein based on its abundance in the bulk supernatant.

Several different SARS-CoV-2 spike ECD constructs were purified in this manner, including full-length, S1, and S2 forms of the wild-type spike (ECD, S1 and S2), the stabilized full-length spike (ECDm) and ECD and ECDm fused to the different trimerization domains. The constructs with the T4-Fib trimerization domain contain StrepTag-II instead of FLAG tags and were purified by IMAC followed by a StrepTrap HP column. As shown in Figure 5, the purified proteins were analyzed by UPLC-SEC (Figure 4A and 4B) with a MALS detector for molecular weight determination (Figure 5C), as well as by SDS-PAGE/Coomassie staining (Figure 5D). As expected, for all of the non-mutated ECD constructs (with and without trimerization domains), the purified protein consists of the S2 subunit with lower levels of S1 and uncleaved full-length protein. By SDS-PAGE, the sizes of S1 and S2 that result from processing of the full-length ECD match well the sizes of S1 and S2 when expressed individually. By UPLC-SEC, purified ECD, S1 and S2 show single predominant peaks of UV absorbance, with lower-abundance, fast-eluting peaks which are likely multimers or aggregates (or uncleaved full-length protein, in the case of ECD). Notably, the MALS molecular weight estimate for the principal peak in the ECD preparation is 75 kDa, which matches the MALS estimate for S2 expressed by itself, supporting the assertion that after purification, the ECD preparation is composed mostly of S2. The mutated ECD (ECDm) is estimated to have a molecular weight of 174 kDa, which is approximately equal to that of the sum of S1 and S2 expressed alone.

**Figure 5:**
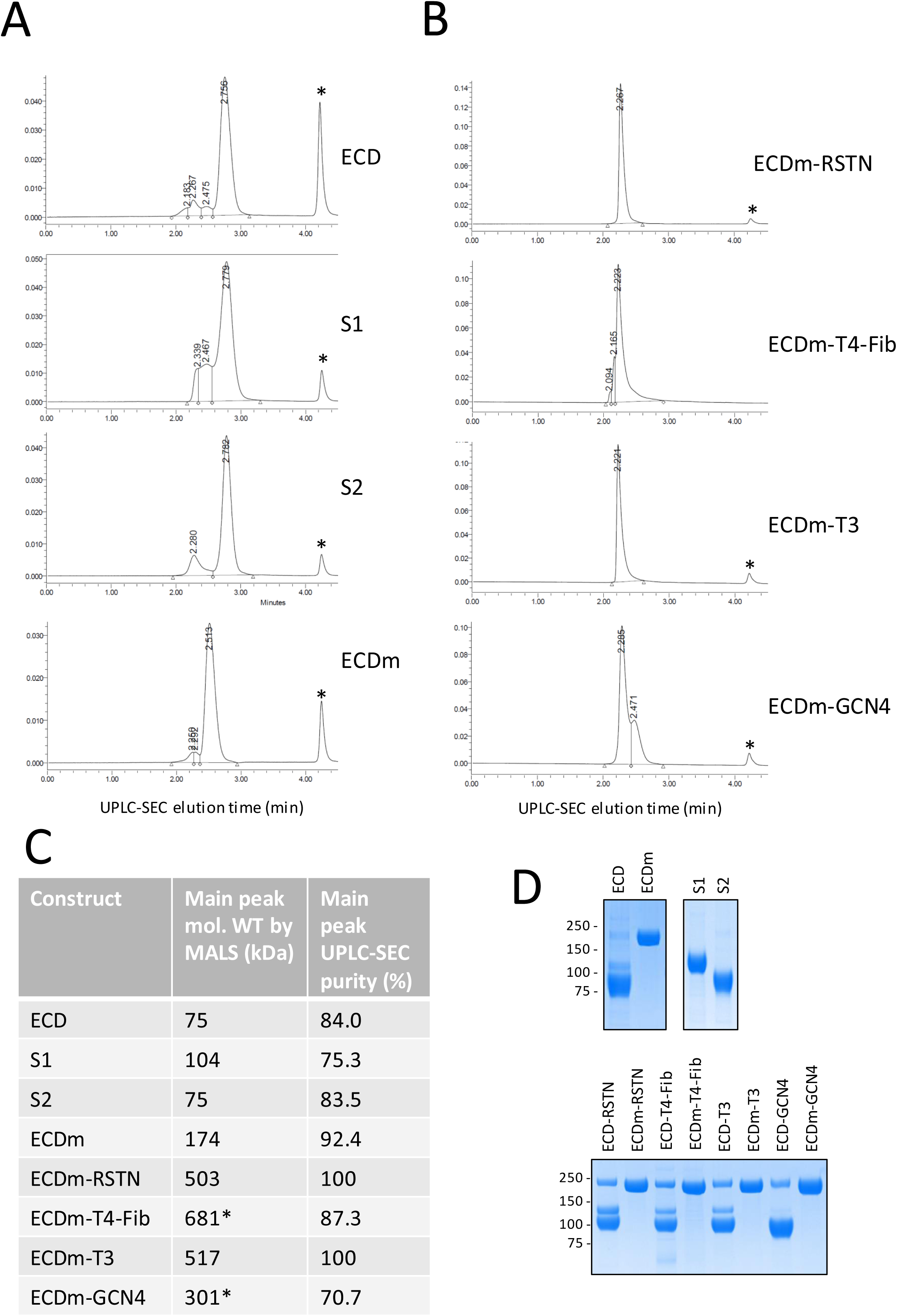
Characterization of IMAC/FLAG-purified spike ectodomain constructs by UPLC-SEC/MALS and SDS-PAGE. (A and B) Purified proteins were separated by UPLC on a BEH200 SEC column. Protein elution was monitored by measuring UV absorbance of eluate (vertical axis). The position of the FLAG peptide elution peak is marked with an asterisk. (C) The molecular weight of the major elution peak for each sample was estimated by MALS. Values marked with asterisks are likely inaccurate due to peak shouldering. (D) 2 μg of each purified protein was analyzed by SDS-PAGE/Coomassie Blue staining.

Purified constructs containing trimerization domains were also analyzed. By SDS-PAGE (Figure 5D, lower panel), they are quite similar, although the amount of S1 lost during purification of the ECD constructs is variable. The mutated (ECDm) trimerization domain fusions all appear to be highly pure by SDS-PAGE. Notably, for the UPLC-SEC analysis, due to varying degrees of loss of S1 from the trimeric ECD constructs, the UPLC-SEC profiles and MALS data are very difficult to interpret (data not shown). For the mutated constructs, the RSTN and T3 fusions give highly uniform purified preparations by UPLC-SEC that are estimated by MALS to have molecular weights of 503 and 517 kDa, respectively. This is consistent with the expected molecular weight of a trimer of ECDm (174 kDa). For the T4-Fib and GCN4 fusions, the principal peaks by UPLC have similar retention times as the other trimerization domains; however, the presence of shoulders on the main peak indicates that it could be composed of more than one species, and therefore, the MALS molecular weight estimates for the trimeric protein are likely inaccurate. Notably, T4-Fib is the most commonly used trimerization domain used in the literature for coronavirus spike proteins. Together, these results indicate that RSTN and T3 fusions are most promising for preparing uniform preparations of SARS-CoV-2 spike ECD with trimeric stoichiometry; additional analyses will be required to evaluate how well these mimic the conformation of the transmembrane spike protein in its native state.

In summary, we have developed a method for transient production in CHO cells of full-length SARS-CoV-2 spike ECD, yielding 100-150 μg/ml within 7 days of plasmid transfection. This is significantly higher than yields reported previously for production of recombinant coronavirus spike proteins. Our results demonstrate that full-length recombinant spike protein is not inherently difficult to produce at high levels in mammalian cells, but that productivity is greatly dependent on host cell line selection: consistent with literature reports, we obtained low yields from transfected HEK293 cells. The mutated construct (ECDm) is likely the best starting point for ongoing development of recombinant protein for therapeutic or diagnostic applications: it can be readily expressed in CHO cells and can be purified to near homogeneity, as assessed by SEC and SDS-PAGE. To mimic the conformation of the spike protein on the virus surface, the RSTN and T3 fusions were most effective at promoting trimeric stoichiometry.

The challenge of manufacturing sufficient amounts of spike protein is likely an important reason why alternative methods, such as mRNA-, adenovirus and AAV-based approaches to induce spike protein expression in vaccinated individuals, have gained prominence with COVID-19 vaccine developers. However, production of vaccine antigen in the form of recombinant spike protein in CHO cells has an intrinsic advantage over these newer technologies: the extensive, decades-long safety record of CHO-derived biologics. The improved yields we have demonstrated in the current study support the feasibility of recombinant, subunit-based vaccines for COVID-19.

Based on past experience with other recombinant proteins, we expected that TGE yields could be further improved in stably transfected CHO pools or clones [21]; indeed, very recently, we have generated stable CHO pools that can reachyields >800 mg/L for ECDm-RSTN (manuscript in preparation). Nonetheless, the speed at which we were able to generate substantial amounts of spike protein by CHO TGE highlights how this approach could complement other methods, in particular for rapid response to health emergencies. As detailed previously [26], we believe that development of alternativesto clone-based CHO production methods, including stable pools and TGE, should be prioritized as potential novel methods for rapid, large-scale manufacturing of biologics.

## 4. ACKNOWLEDGEMENTS

We thank the Quality Attributes and Characterization Team at the National Research Council of Canada for performing UPLC-SEC/MALS analyses.

## SUPPLEMENTARY MATERIAL CAPTIONS

**Supplementary Figure 1:**
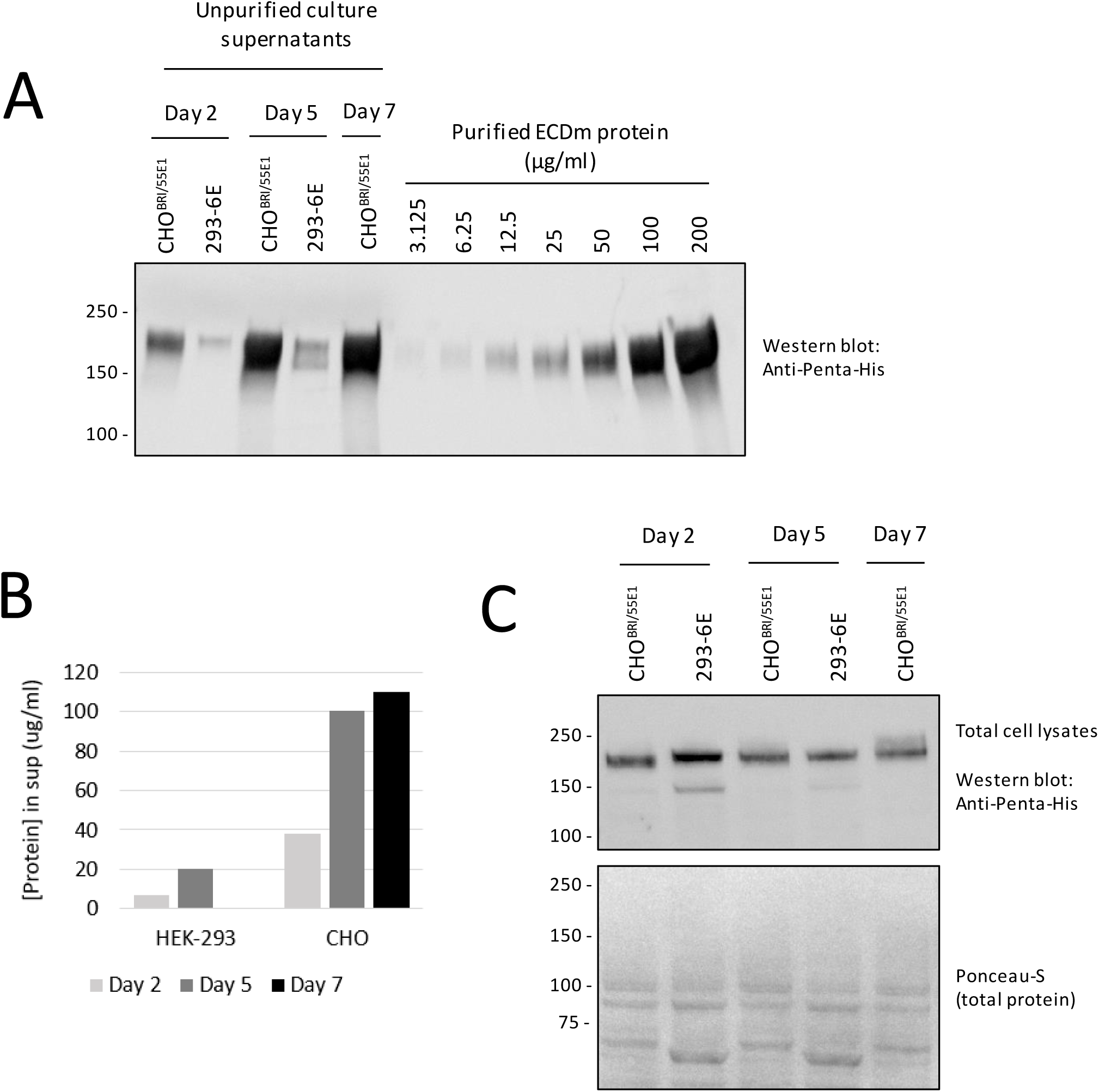
Recombinant ECD concentrations in culture supernatants are higher in CHO vs. HEK293 cells, but intracellular protein levels are similar. CHO^BRI/55E1^ and 293-6E cells were transfected with the ECDm expression plasmid following standard protocols. Samples were taken at 2, 5 or 7 days post-transfection for preparation of cleared culture supernatants and cell pellets (for lysis and total protein extraction). (A) Cleared supernatants (10 μl) were analyzed by western blotting along with known amounts of IMAC/FLAG-purified ECDm protein. Densitometry analysis of the purified protein bands was used to prepare a standard curve and estimate ECD protein concentrations in the supernatants (B). (C) Western blot of cell lysates for detection of intracellular ECD protein. Protein lysates were prepared from cell pellets and protein concentrations were measured; equal amounts of total protein were analyzed by western blotting.

